# 5′ UTR length regulates alternative N-terminal protein isoform production in health and disease

**DOI:** 10.64898/2026.01.20.700518

**Authors:** Jimmy Ly, Eric M. Smith, Matteo Di Bernardo, Yi Fei Tao, Elizabeth M. Black, Ekaterina Khalizeva, Iain M Cheeseman

## Abstract

The 5′ untranslated region (5′ UTR) of an mRNA is classically viewed as a regulatory region that controls the amount of protein production, but not the resulting protein sequence. Here, we demonstrate that 5′ UTR length plays a direct role in alternative N-terminal protein isoform production by controlling start codon selection. We find that very short 5′ UTRs enhance leaky ribosome scanning, thereby promoting the production of truncated alternative N-terminal protein isoforms. We also show that endogenous changes in 5′ UTR length due to alternative transcription initiation can tune the relative abundance of alternative N-terminal isoforms from the same gene. In addition, we identify mutations in rare genetic diseases that alter 5′ UTR length, including a deletion in the VHL 5′ UTR in von Hippel–Lindau disease that shifts translation toward the shorter VHLp19 isoform. Together, our results implicate 5′ UTR length as a determinant of alternative N-terminal isoform production and reveal an underappreciated mechanism by which noncoding changes can reshape the proteome.

**Highlights:**

- 5′ UTR length affects the landscape of endogenous alternative N-terminal protein isoforms
- Generation of an alternative truncated AKR7A2 isoform is mediated by short 5′ UTR length
- Alternative transcription initiation modulates 5′ UTR length to tune N-terminal isoform ratios
- Pathogenic VHL 5′ UTR variants perturb N-terminal isoform ratios by altering 5′ UTR length

## Introduction

For decades, eukaryotic mRNAs were generally assumed to encode a single protein product. This view has been revised by growing evidence that many transcripts initiate translation at multiple, in-frame alternative start codons, producing distinct N-terminal protein isoforms from a single mRNA (*1–6*). These alternative N-terminal isoforms can differ from their annotated counterparts in their interacting partners (*7–9*), subcellular localization (*10–13*), stability (*14–16*), and other fundamental aspects of protein biology (*17, 18*). Alternative translation initiation substantially expands proteomic diversity beyond current annotations and contributes broadly to cellular physiology and disease. Defining the mechanisms that govern the production of alternative N-terminal isoforms is therefore essential for understanding this previously hidden layer of gene expression.

Eukaryotic mRNAs are typically organized into three functional regions: the protein-coding open reading frame (ORF), an upstream 5′ untranslated region (5′ UTR), and a downstream 3′ untranslated region (3′ UTR), with the 5′ UTR playing key roles in regulating translational efficiency (*19–21*). Numerous cis-regulatory features within the 5′ UTR can influence start codon selection (*22–25*), including the Kozak context for the start codon (*26–30*), the use of non-AUG initiation codons (*1, 2, 6, 28*), RNA secondary structure (*11, 31–33*), and upstream open reading frames (uORFs) (*24, 34, 35*). In contrast, the length of the 5′ UTR itself has not traditionally been considered a major determinant of the corresponding protein products that are produced, including alternative N-terminal isoform production. Prior work by Marilyn Kozak demonstrated that shortening the 5′ UTR of a reporter mRNA to below 30 nucleotides (nt) promotes leaky ribosomal scanning (*36, 37*). In addition, recent structural studies support a model in which the ribosome engages the 5′ end of the mRNA in a manner that occludes an 40-50 nt region proximal to the cap from undergoing robust initiation (*38*). Consequently, start codons positioned very close to the 5′ cap may be inefficiently recognized or bypassed completely. Although these findings suggest 5′ UTR length could act as an important determinant of start codon selection *in vitro*, the extent to which endogenous transcripts exploit this mechanism to produce alternative N-terminal isoforms remains largely unexplored.

Here, we use ribosome profiling (*5, 28*) and Cap Analysis of Gene Expression (CAGE)-sequencing (*39*) to identify endogenous mRNAs that undergo short-5′-UTR-driven leaky scanning to produce functional alternative N-terminal isoforms. We further show that alternative transcription initiation can modulate 5′ UTR length, thereby altering protein isoform production without changing the coding sequence. Finally, we identify pathogenic patient alleles that alter 5′ UTR length and consequently shift alternative N-terminal isoform ratios, highlighting the physiological importance of 5′ UTR lengths.

## Results

### 5′ UTR length regulates alternative N-terminal isoform selection for AKR7A2

Our recent work using start site ribosome profiling (*28*) identified hundreds of human genes that encode dual protein products with distinct subcellular localizations (*10*). Importantly, a subset of these mRNAs exhibits alternative start codon usage that cannot be readily explained by known translational control mechanisms. One such example is AKR7A2 (*40*), which contains two in-frame translation start sites as revealed by start site ribosome profiling (Fig. 1A). Translation from the annotated upstream AUG generates a mitochondrial protein, whereas initiation at a downstream AUG produces a cytosolic/nuclear isoform (Fig. EV1A; (*10*)). To determine how these AKR7A2 isoforms are decoded, we expressed an AKR7A2 construct containing its native 5′ UTR and complete coding sequence fused to GFP. When expressed from a dox-inducible promoter (P_TRE3G_; (*41, 42*)), this 5′ UTR-AKR7A1-GFP construct produced exclusively mitochondrial localization, indicating strict initiation at the first AUG (Fig. 1B). Although the upstream AUG has a moderate strength Kozak context (Fig. 1A), this result suggests that Kozak strength alone is insufficient to drive leaky scanning for this transcript to allow downstream initiation. In contrast, transfection of an *in vitro*–transcribed AKR7A2 5′ UTR–CDS–GFP mRNA resulted in clear dual localization to mitochondria, cytosol, and nucleus (Fig. 1B), consistent with usage of both start codons and the behavior of endogenous AKR7A2.

**Figure 1.**
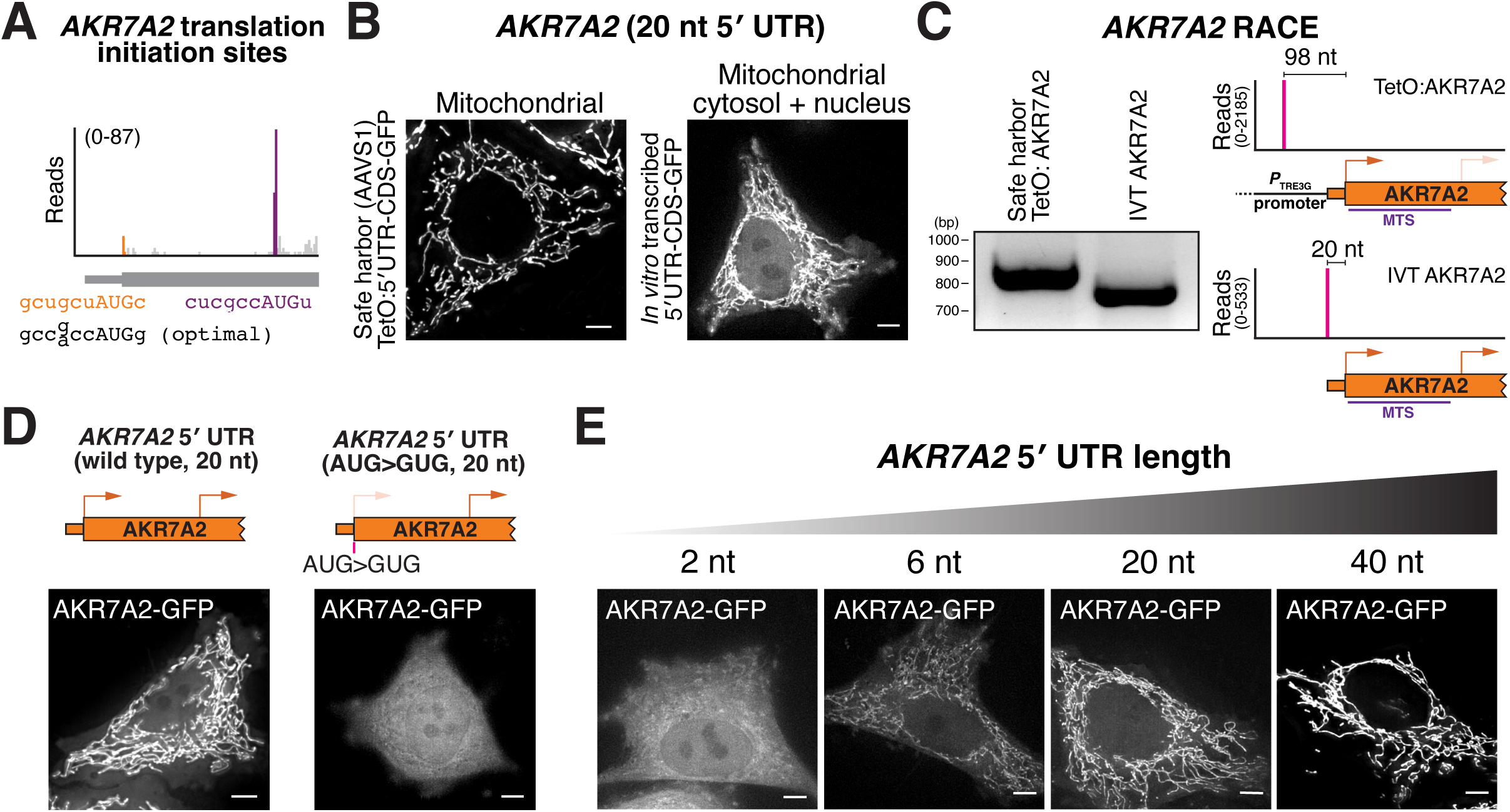
5′ UTR length regulates AKR7A2 alternative N-terminal isoform selection. (**A**) Read distribution from translation initiation site profiling around the AKR7A2 alternative start codons. The orange peak represents reads at the annotated start codon whereas the purple reads represent the alternative N-terminally truncated start site. Optimal Kozak context from (*26*). (**B**) Live-cell imaging of AKR7A2-GFP produced from the dox-inducible promoter or in vitro transcribed mRNA. (**C**) Left, 5′ RACE of endogenous AKR7A2 from transgenic AKR7A2 expressed under dox-inducible promoter and in vitro transcribed AKR7A2 mRNA. Right, quantification of 5′ RACE reads by sequencing. (**D**) Left, live cell imaging of AKR7A2 after transfecting mRNA containing the AKR7A2 5′ UTR – CDS – GFP. Right, localization of AKR7A2 where the annotated start codon was mutated to GUG. (**E**) Live cell imaging of cells transfected with in vitro transcribed AKR7A2 5′ UTR – CDS – GFP with the indicated 5′ UTR lengths. Scale bar represents 5 µm

To understand the difference in AKR7A2 localization produced from the dox-inducible promoter vs. the *in vitro* transcribed mRNA, we considered the nature of these constructs. Most promoters used for ectopic expression in human cells will append additional nucleotides upstream of the cloned 5′ UTR to the transgenic mRNA (*43–45*). For example, the dox-inducible promoter (*P*_TRE3G_) used for our experiments will add ∼78 nucleotides to the cloned mRNA sequence (Fig. 1C). Thus, the dox-inducible mRNA has a longer 5′ UTR than the *in vitro* transcribed AKR7A2 construct.

The native AKR7A2 5′ UTR is unusually short, only 20 nt, whereas the average 5′ UTR length in humans is around 200 nucleotides (*19*). To test whether 5′ UTR length underlies these observed differences alternative N-terminal isoform production, we transfected cells with *in vitro* transcribed AKR7A2 mRNAs containing defined 5′ UTRs lengths and a C-terminal GFP tag. The native 20-nt AKR7A2 5′ UTR supported production of both mitochondrial and cytosolic/nuclear isoforms, whereas mutation of the first start codon (AUG to GUG) abolished mitochondrial localization, confirming that relative localization reflects initiation at the two start codons (Fig. 1D). Extending the 5′ UTR to 40 nt using the Xenopus β-globin sequence suppressed downstream initiation and yielded exclusively mitochondrial protein localization (Fig. 1E). Conversely, further shortening the 5′ UTR to 6 or 2 nt shifted initiation toward the downstream start site, favoring cytosolic/nuclear localization (Fig.1E). Notably, mouse AKR7A2 also has an unusually short 11-nt 5′ UTR and produces dual-localized protein isoforms in a manner that depends on 5′ UTR length (Fig. EV1B-D). However, 5′ UTR length is not universally conserved as chimpanzee and gorilla AKR7A2 transcripts have substantially longer 5′ UTRs (67 and 127 nt, respectively; (Fig. EV1B). Whether these differences alter leaky scanning and isoform production across species or are errors in transcript annotations remains to be determined. Overall, these results demonstrate that the short 5′ UTR length of AKR7A2 strongly influences the production of its differentially localized alternative N-terminal isoforms.

### Nuclear-encoded mitochondrial mRNAs are enriched for short 5′ UTRs

Based on our analysis of AKR7A2, we hypothesized that other mRNAs with short 5′ UTRs would be more likely to produce N-terminally truncated protein isoforms. To evaluate whether short 5′ UTR lengths are correlated with the production of alternative translational isoforms on a global level, we conducted a computational analysis of the relationship between 5′ UTR length (based on annotated mRNAs; (*46*)) and downstream translation initiation (based on start site ribosome profiling; (*28*)). We found that mRNAs with short 5′ UTRs (≤40 nt) produced a larger frequency of alternative N-terminally truncated isoforms compared to those mRNAs with longer 5′ UTRs (>40 nt) (Fig. 2B; Fig. EV2A-B, Table S1). Amongst diverse biological processes, we found that nuclear-encoded genes that produce mitochondrial-localized proteins, such as AKR7A2, are enriched for short 5′ UTRs (Fig 2B-C; Fig. EV2C-E).

**Figure 2.**
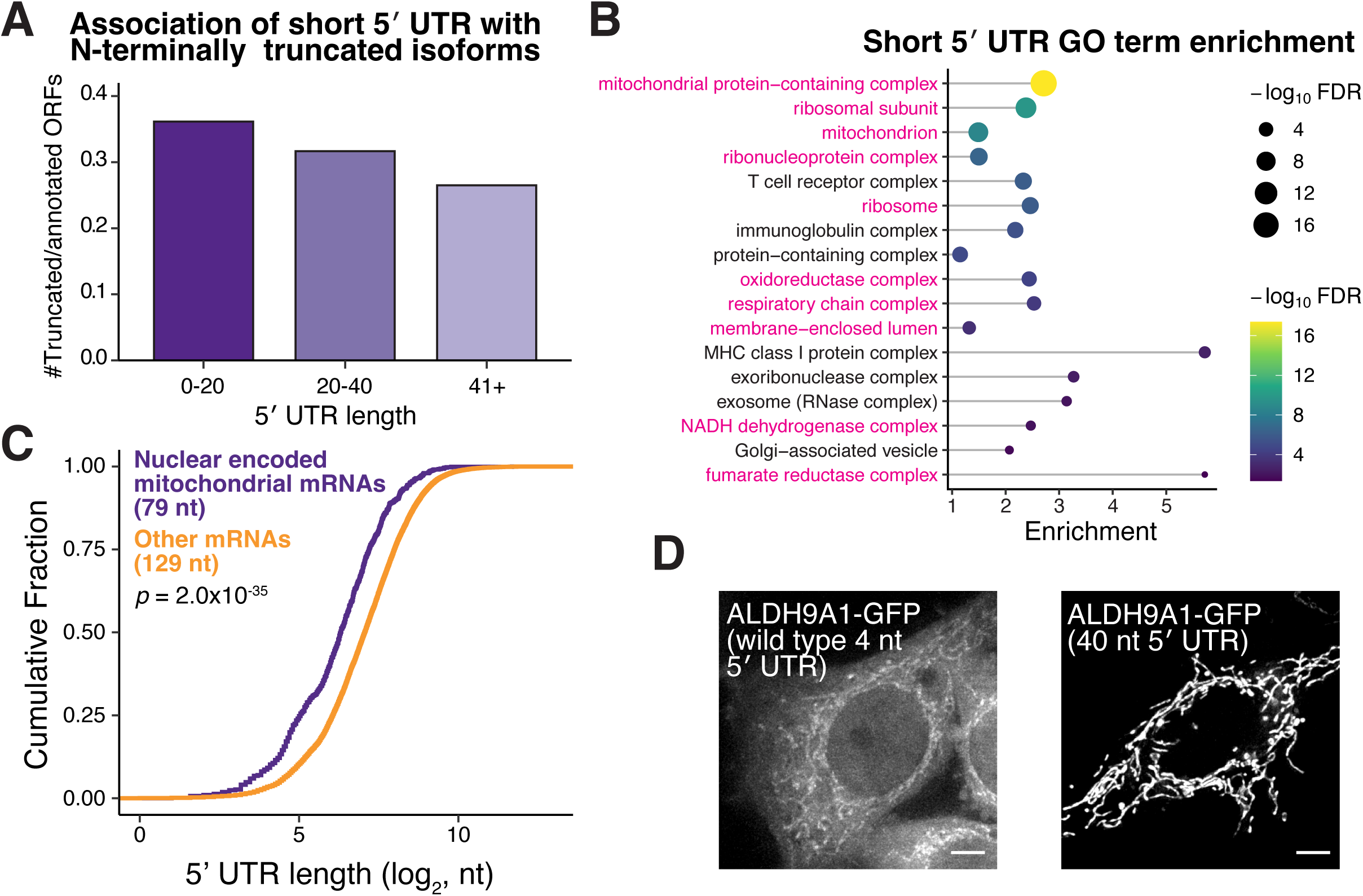
Short 5′ UTR regulation affects many mitochondrial genes. (**A**) Bar plot showing the relative proportion of N-terminal truncations to annotated ORFs for genes in different 5′ UTR length bins indicating that mRNAs with shorter 5′ UTRs have more N-terminal truncations. (**B**) GO term analysis of human genes with short 5′ UTRs. Pink GO terms are highlighted in magenta. (**C**) Cumulative distribution fraction plots showing the 5′ UTR lengths of nuclear-encoded mitochondrial genes compared to all other nuclear-encoded genes. Statistic represents results from Wilcoxon rank sum test. (**D**) Live cell imaging of cells transfected with in vitro transcribed ALDH9A1 5′ UTR – CDS – GFP with the wild type (4 nt) and elongated (40 nt) 5′ UTR. Scale bar represents 5 µm.

Based on our analysis, we identified at least 75 human genes with short 5′ UTRs that produce more than one translational isoform based on start site ribosome profiling (Table S1). To test these, we focused on ALDH9A1 (*47, 48*), which has an annotated 5′ UTR length of 4 nt (Fig. EV2D). Indeed, using mRNA transfection, the annotated ALDH9A1 mRNA produced dual-localized mitochondrial and cytosolic isoforms (Fig. 2D). In contrast, increasing the 5′ UTR length to 40 nt biased translation towards the mitochondrial isoform whereas reducing the length biased towards the production of the cytosolic isoform (Fig. 2D; Fig. EV3A-B). We next examined the conservation of the short ALDH9A1 5′ UTR length. Analysis of ALDH9A1 5′ UTR lengths across mammals based on the Ensembl database (*46*) revealed that the short 5′ UTR is not highly conserved (Fig. EV3C). Despite this, orangutan ALDH9A1 mRNA, which contains a 40-nt 5′ UTR, still produced dual-localized ALDH9A1 protein isoforms (Fig. EV3D-E). Analysis of the first AUG start codon for orangutan ALDH9A1 revealed a weak Kozak context, which may promote the production of dual-localized ALDH9A1 isoforms (Fig. EV3F). Much like with AKR7A2, these results suggest an evolutionary pressure to maintain the production of dual ALDH9A1 translational isoforms, but that this can occur through distinct molecular and regulatory mechanisms.

Overall, these data suggest that 5′ UTR length regulation can act as a broad paradigm to affect the translation of alternate protein isoforms, contributing to diverse biological processes and expanding the functional human proteome, with a particular role in mitochondrial function.

### Alternative transcription regulates start codon selection of AKR7A2 and Complement Factor D (*CFD*)

Given the importance of 5′ UTR length in dictating protein isoform selection, we next considered mechanisms that could modulate 5′ UTR length to influence start codon selection. One such mechanism is alternative transcription initiation, which can generate mRNAs with distinct 5′ UTRs. Alternative transcription initiation in budding yeast has been shown to influence the production of alternative N-terminal protein isoforms by changing the 5′ end of the mRNA to include or exclude start codons (*12*). To identify analogous cases in human cells, we performed Cap Analysis of Gene Expression (CAGE; (*39*)) in HeLa cells to identify alternative transcription start sites (Fig. 3A). In total, we identified 81079 transcription start sites, with 86% (7714/8314) of genes displaying more than one start site (Table S2). Of genes with more than one transcription start site, our analysis revealed 8468 alternative transcription initiation sites that are downstream of the annotated start codon and are thus predicted to cause the production of a truncated protein isoform (Table S2). For example, AKR7A2 has a downstream promoter that excludes the annotated start codon in the mRNA such that this transcript fails to translate the mitochondrial isoform (Fig. EV4A). As a result, transcripts initiated from this alternative promoter can only produce the truncated nuclear/cytosolic isoform. Thus, AKR7A2 uses multiple mechanisms to generate the alternative truncated protein isoform, including short 5′ UTR–driven leaky scanning from the annotated mRNA isoform (ENST00000235835) as well as alternative promoter usage.

**Figure 3.**
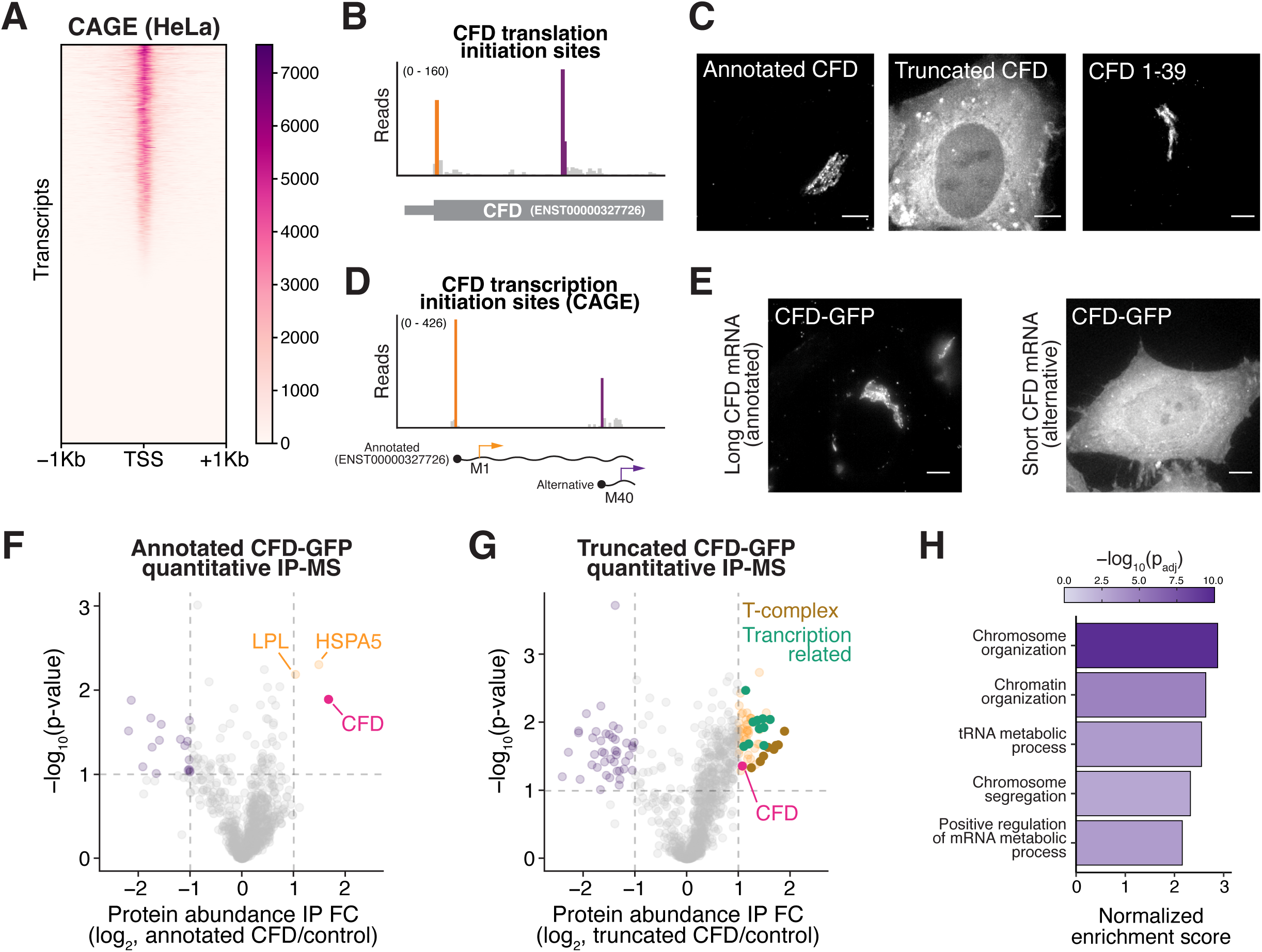
Alternative transcription of CFD produces an intracellular alternative N-terminal isoform. (**A**) Heatmap of CAGE-Seq data from HeLa cells. Scale bar represents scaled read counts. (**B**) Read distribution trace from translation initiation site profiling around the CFD alternative start codons. (**C**) Live cell imaging of annotated CFD, Truncated CFD isoforms and the N-terminus of annotated CFD tagged with a C-terminal GFP. Images are not scaled equally. (**D**) Read distribution traces from CAGE-Seq data around the CFD transcription start sites. (**E**) Live cell imaging of HeLa cells expressing the annotated CFD mRNA isoform (left) or the alternative transcript isoform (right). Images are not scaled equally. Scale bar represents 5 µm. (**F**) Volcano plot of quantitative IP-MS analysis of annotated CFD. (**G**) Volcano plot of quantitative IP-MS analysis of truncated CFD. n = 2 biological replicates. (**H**) Gene set enrichment analysis for truncated CFD interactors.

A second compelling example of alternate promoters affecting translation start site usage is the Complement Factor D (CFD) gene (*49*), which contains two conserved translation initiation sites (Fig. 3B; Fig. EV4B). These sites are predicted to generate alternative N-terminal isoforms with distinct subcellular localizations (Fig. EV4C-D). To evaluate this, we tested the localization of each isoform in HeLa cells. Indeed, translation from the annotated start codon produced a Golgi-localized protein (Fig. 3C), consistent with CFD’s role as a secreted complement factor (*49*). In contrast, initiation at the downstream site produced a truncated isoform that localized to the cytosol and nucleus (Fig. 3C). Notably, we found that expression of the annotated CFD mRNA produces only the Golgi-localized isoform (Fig. 3D-E), indicating that this transcript does not undergo leaky ribosomal scanning. Instead, 5′ CAGE analysis revealed an alternative downstream transcription start site (Fig. 3D) that removes the annotated start codon and enforces production of the truncated CFD isoform (Fig. 3E).

The alternative start codon and differential localization for CFD isoforms are highly conserved across vertebrates (Fig. EV4E). However, a functional role for the intracellular CFD isoform has yet to be described. To test this, we next investigated the role of this truncated CFD isoform. Based on immunoprecipitations coupled with quantitative mass spectrometry, the annotated, secreted CFD isoform interacted with only two ER/Golgi-associated proteins, consistent with its localization to the secretory pathway (Fig. 3F). In contrast, the truncated isoform showed a broad interaction network, including numerous proteins involved in genome organization and transcription (Fig. 3G-H; Table S3). CFD is not essential for viability in cultured human cells (DepMap; (*50*)), limiting functional analysis of the intracellular isoform in cultured cell lines. Nevertheless, ClinVar (*51*) reports a mutation (ClinVar accessions: VCV001511411.6 and VCV001494493.5) in the alternative CFD start codon that is predicted to disrupt the translation of intracellular CFD. This suggests that the alternative CFD isoform may perform physiologically important intracellular function, with differential promoter usage affecting the relative production of these isoforms across conditions.

Overall, these data highlight the importance of promoter selection in regulating the production of alternative N-terminal isoforms.

### Alternative transcription can control 5′ UTR length to regulate isoform selection of ALDH9A1 and GUK1

The examples above illustrate how substantial differences in alternative transcription start sites can remove upstream start codons, redirecting translation to downstream sites. Importantly, our analysis of AKR7A2 and ALDH9A1 (Fig. 1 and 2) highlighted that small changes in 5′ UTR length alone—without altering coding sequences—can also tune protein isoform production. To test whether alternative transcription can change the production of alternative N-terminal isoforms by modulating 5′ UTR length, we analyzed our CAGE-seq data to identify alternative promoters that strictly change 5′ UTR length (Table S2). For example, the annotated ALDH9A1 transcript has a 4 nt 5′ UTR, but alternative transcription initiation extends the 5′ UTR to 28 nt (Fig. 4A). Notably, we found that the longer 28 nt 5′ UTR isoform increased initiation at the first start codon elevating the relative levels of the mitochondrial isoform (Fig. 4B-C).

**Figure 4.**
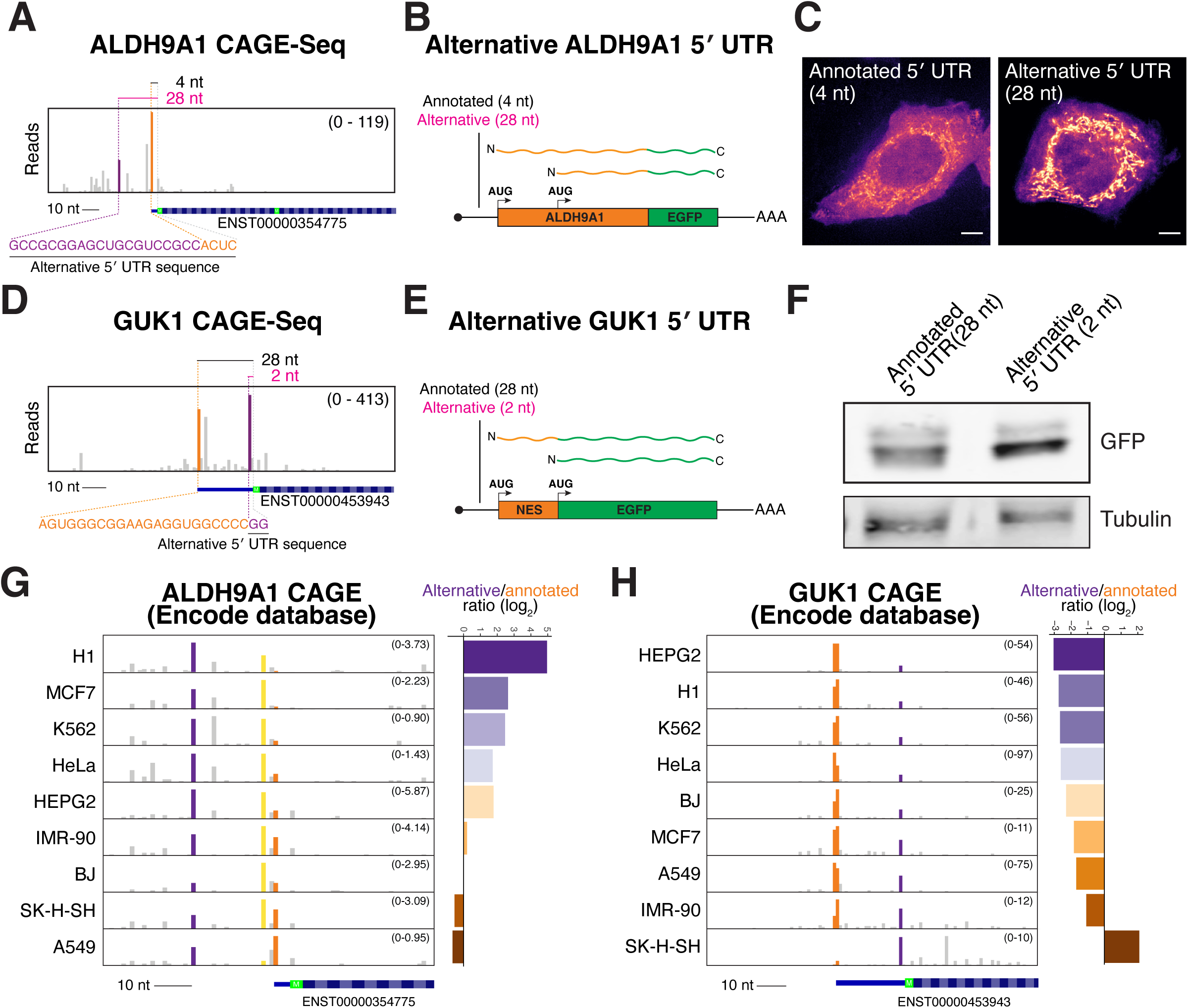
Alternative transcription initiation modulates 5′ UTR length to regulate alternative N-terminal isoform ratios. (**A**) CAGE-Seq analysis of ALDH9A1 transcription start sites. Orange reads represent the annotated transcription start site and purple reads represent the alternative transcription start site resulting in a longer 5′ UTR. (**B**) Schematic of in vitro transcribed ALDH9A1 reporter mRNA with annotated or alternative 5′ UTRs. The relative ratio of mitochondrial compared to nuclear/cytosolic isoform represents the relative start codon usage. (**C**) Live cell imaging for wild type (4 nt) and alternative (28 nt) ALDH9A1 reporters. Scale bar indicates 5 µm. (**D**) Same as (A) except for GUK1. (**E**) Schematic of in vitro transcribed GUK1 5′ UTR reporters. The GUK1 5′ UTRs were appended to a nuclear export signal flanked with AUG start codons followed by a GFP. (**F**) Western blot showing the relative ratios of the protein products produced by the first start codon (larger protein) vs the second start codon (shorter protein). (**G**) CAGE-seq trace for ALDH9A1 transcription start sites across 9 cell lines from the ENCODE consortium (*55*). The orange read indicates reads at the annotated transcription start site and the purple indicates alternative longer 5′ UTR isoform. The yellow reads indicate another alternative start site, but only extending it to 7 nt. The quantification on the right indicates the relative number of alternative promoter reads normalized to the annotated start site. (**H**) Same as (G) except with GUK1.

Reciprocally, we identified alternative transcription start sites that shorten the 5′ UTR relative to the annotated transcript. GUK1 (*52, 53*), which contains two translation initiation sites (Fig. EV5A), has an annotated 5′ UTR of 28 nt. However, usage of an alternative transcription start site generates an mRNA isoform with a 2 nt 5′ UTR without changing the coding sequence (Fig. 4D). Because the alternate translation initiation sites in GUK1 do not produce differentially localized protein isoforms (Fig. EV5B), we generated a reporter construct to assess the usage of these two start sites (Fig. 4E). This reporter construct contains two in frame AUG start codons with an intervening nuclear export signal followed by a C-terminal GFP (Fig. 4E). Usage of the different start codons will produce protein products of different sizes such that the relative ratio of the proteins will reflect the relative extent of leaky ribosome scanning. By comparing the amount of leaky scanning for the annotated GUK1 5′ UTR (28 nt) relative to the alternative GUK1 5′ UTR (2 nt), we observed enhanced leaky ribosome scanning for the 2 nt 5′ UTR (Fig. 4F; Fig. EV5C). Therefore, despite encoding identical open reading frames, endogenous transcripts with differing 5′ UTR lengths will produce distinct ratios of GUK1 protein isoforms. These data highlight the importance of considering alternative transcripts with different 5′ UTR lengths when interpreting the protein products generated from a given gene.

Our prior work identified cell type–specific alternative transcription of KNSTRN/SKAP, which results in the production of a distinct, testes-specific N-terminal isoform (*54*). Thus, changes in 5’ UTR length could result in differing tissue or context-specific production of alternate protein isoforms. To test this, we examined whether alternative transcription initiation of ALDH9A1 and GUK1 varies across biological contexts thereby modulating alternative N-terminal isoform production solely by changing 5′ UTR length. Based on analysis CAGE-seq datasets generated by the ENCODE consortium (*55*), we identified diverse promoter usage across diverse cancer cell lines. For both ALDH9A1 and GUK1, transcription start site usage varies markedly across a panel of 9 cancer cell lines (Fig. 4G-H), with the ratios of longer to shorter 5′ UTR transcripts spanning 48-fold for ALDH9A1 and 31-fold for GUK1. These results indicate that alternative transcription initiation of ALDH9A1 and GUK1 differs across cell types and may function to tune N-terminal isoform abundance in a context-specific manner.

### Pathogenic 5′ UTR mutations regulate alternative start codon selection through 5′ UTR length control

The analysis of rare human disease typically focuses on coding mutations (*56–58*). However, our analysis of 5′ UTR lengths suggests that non-coding changes could also alter relative isoform production. Thus, we considered whether there are pathological mutations that alter 5′ UTR length and start codon selection. To test this, we mined the ClinVar database (*51*) for rare disease alleles that are predicted to alter 5′ UTR length. In total, we identified 988 alleles across 519 genes that are predicted to change 5′ UTR length (Fig. 5A; Table S4). Most variants were small insertions, duplications, or deletions of one or two nucleotides. However, we identified several variants that are predicted to cause substantial alterations in 5′ UTR length ranging from −411 nt to +889 nt (Fig. 5A).

**Figure 5.**
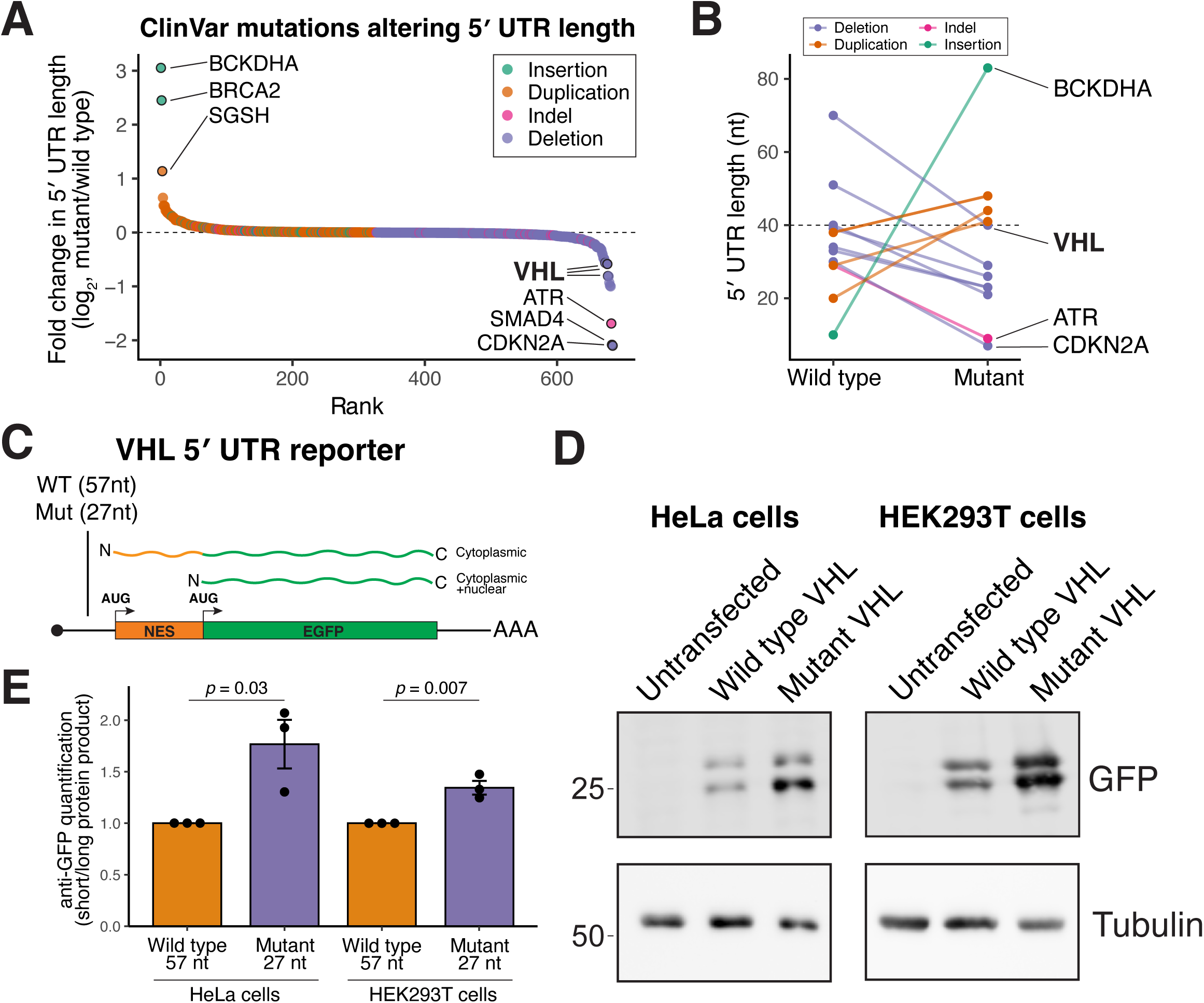
5′ UTR disease mutations can alter alternative N-terminal isoform ratios. (**A**) Water fall plot showing the fold change in 5′ UTR length induced by ClinVar mutations. (**B**) Subset of ClinVar 5′ UTR mutations that alter length within the 5′ UTR range of 40 nt. (**C**) Schematic of VHL 5′ UTR reporter. The wild type and mutant VHL 5′ UTR is driving the expression of nuclear export signal flanked by two start codons followed by a GFP. The first start codon produces a NES-GFP that should be enriched in the cytosol, whereas the second start produces just GFP which should localize to both the nucleus and cytosol. (**D**) Representative Western blot from HeLa (left) or HEK293 (right) cells transfected with the indicated in vitro transcribed VHL reporters. (**E**) Quantification of western blots in D. Error bar indicates standard error of the mean, n = 3 biological replicates, and statistics indicate student’s T-test.

We next focused on variants predicted to have the strongest impact on start codon selection by affecting transcripts with very short 5′ UTRs (<40 nt). We identified multiple ClinVar mutations that are predicted to increase or decrease short UTR by at least 10 nt (Fig. 5B). We identified 5 mutations that lengthen short 5′ UTRs, which we predict would suppress downstream leaky ribosome scanning, and 5 mutations that shorten these already short 5′ UTRs, which is predicted to enhance leaky scanning (Table S4). In addition, we identified cases in which pathogenic deletions convert long 5′ UTRs into short (<40 nt) UTRs, a change expected to promote leaky ribosome scanning. Using these criteria, we identified CRYGD, MYD88, and VHL as candidate genes in which disease-associated variants may induce or enhance leaky ribosome scanning resulting in a N-terminally truncated protein isoform.

To test the impact of these 5′ UTR changes on alternative N-terminal isoform selection, we focused on VHL. VHL is a tumor suppressor that functions as part of an E3 ubiquitin ligase complex that targets hypoxia-inducible factor (HIF) for ubiquitination and degradation (*59*). Mutations in VHL cause von Hippel–Lindau disease, a hereditary cancer syndrome (*60*). VHL is alternatively translated to produce ∼30 kDa and ∼19 kDa isoforms (*61, 62*), with unique functions (*63–66*). Notably, mutations that affect the ratio of alternative VHL isoforms differentially contribute to disease and pathology (*67, 68*).

The annotated VHL mRNA contains a 5′ UTR of 70 nt. However, empirical mapping of transcription start sites by CAGE revealed that the predominant VHL 5′ UTR in HeLa cells is shorter, at 57 nt (Fig. EV6A). We therefore used the GFP reporter described above (Fig. 5E) to assess start codon selection driven by either the wild-type VHL 5′ UTR (57 nt) or a pathogenic 30 nt 5′ UTR deletion mutant (VCV002928001.4) (Fig. 6C). The 5′ UTR deletion enhanced leaky ribosome scanning (Fig. 6D-E; Fig. EV6B-C) and is therefore predicted to increase production of the shorter VHL p19 isoform. Consistent with this prediction, analysis of GFP reporter localization also supported increased leaky scanning in the presence of the pathogenic 5′ UTR deletion. Together, these results demonstrate that disease-associated 5′ UTR mutations can directly regulate alternative start codon usage, altering N-terminal isoform output and highlighting the importance of evaluating 5′ UTR variants beyond their canonical regulatory roles. As non-coding variants are typically underreported in rare disease databases (*56, 69*), the prevalence of such alleles may be an under representation of potential genetic impacts across human disease.

**Figure 6.**
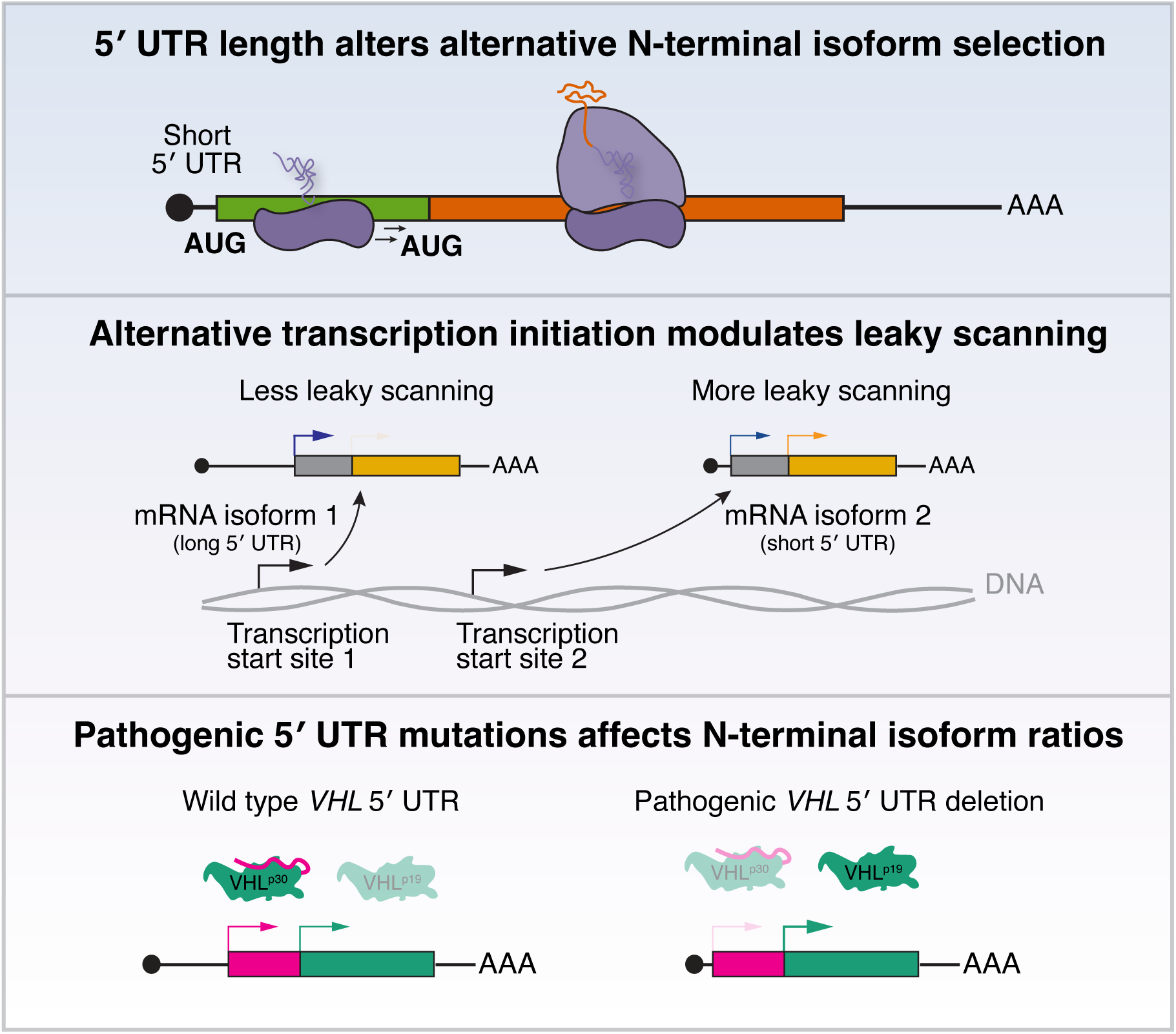
Dynamic changes in 5′ UTR regulates alternative N-terminal isoform production. Schematic model for how changes in 5′ UTR length can tune alternative N-terminal isoform ratios in health and disease.

## Discussion

Defining the cis-regulatory features that control the relative production of alternative N-terminal isoform in mammals has important relevance to core cellular function and disease. Here, we demonstrate that endogenous 5′ UTR length is an important determinant that tunes the relative usage of alternative start codons (Fig. 6). For example, both AKR7A2 and ALDH9A1 mRNAs undergo leaky ribosomal scanning to generate distinct mitochondrial and nuclear/cytosolic N-terminal isoforms. Notably, these transcripts have exceptionally short 5′ UTRs (20 nt for AKR7A2 and 4 nt for ALDH9A1) and extending the 5′ UTR length suppresses leaky scanning in a graded manner, promoting the production of their longer, mitochondrial isoforms (Fig. 1-2).

The importance of 5′ UTR length in regulating alternative isoform production has practical implications for transgenic expression systems, as commonly used promoters often append additional sequences to the 5′ end of transcripts. For example, the TRE3G promoter adds 78 nt to the 5′ UTR (Fig. 1C), abolishing leaky scanning of AKR7A2 and ALDH9A1. Thus, studies of alternative start codon usage must carefully consider reporter design, since plasmid- or transgene-based systems can inadvertently alter 5′ UTR architecture or introduce confounding effects on transcription and splicing (*70*). In such cases, *in vitro*–transcribed mRNA reporters with defined 5′ ends may provide a more accurate approach for assessing start codon selection.

Based on existing genome annotations, there are 2899 genes with at least one mRNA isoform containing a short 5′ UTR (Table S1). However, because not all of these transcripts are expressed in HeLa cells—the cell line used for our ribosome profiling analysis (*28*)—we cannot directly assess whether each of these genes also use alternative translation start sites. Nonetheless, given their short 5′ UTR lengths, the potential for alternative translation initiation should be considered when analyzing these genes.

We further show that alternative promoter usage can alter 5′ UTR length, and that even modest changes are sufficient to shift the relative production of N-terminal protein isoforms. For example, the annotated GUK1 transcript contains a 28-nt 5′ UTR, whereas an alternative transcription start site generates an mRNA with a 2-nt 5′ UTR, promoting leaky ribosome scanning and altered start codon selection. Thus, endogenous changes in promoter usage can tune start codon selection efficiency and reshape protein isoform output. Notably, this alternative GUK1 transcript is absent from current genome annotations, underscoring how incomplete transcript models obscure proteomic diversity. Finally, by analyzing the ClinVar database (*51*), we identified disease-associated alleles predicted to change 5′ UTR length (Fig. 5). Strikingly, 10 of these alleles fall within a short-UTR regime—either occurring in transcripts with 5′ UTRs shorter than 40 nt or truncating longer 5′ UTRs to below this 40 nt threshold—where leaky scanning is predicted to be induced. Among these, we identified pathogenic variants in the VHL 5′ UTR linked to Von Hippel–Lindau syndrome that markedly shorten the 5′ UTR (Fig. 5). We show that these mutations promote leaky ribosome scanning, thereby shifting translation toward a truncated VHL isoform. Together, these findings establish altered 5′ UTR length as a previously underappreciated disease mechanism, demonstrating that noncoding mutations can drive pathology not by altering mRNA stability or overall translational efficiency, but by rewiring start codon selection and protein isoform output. As such, 5′ UTR mutations represent a critical and often overlooked class of pathogenic variants that should be systematically considered in human genetic disease.

## Data availability

The CAGE-Seq data was deposited to GEO with the identifier GSE316181. The mass spectrometry proteomics data have been deposited to the ProteomeXchange Consortium via the PRIDE (*71*) partner repository with the dataset identifier PXD073007 and 10.6019/PXD073007. Code and analysis of 5′ UTR ClinVar variants are available at https://github.com/cheeseman-lab/5-utr-clinvar-mutations.

## Acknowledgements

This work was supported by a grant from the NIH (R35GM126930 to I.M.C) and the Chan Zuckerberg Initiative Rare as One Project grant to I.M.C. J.L. and Y.F.T. are supported in part by the Natural Sciences and Engineering Research Council of Canada. E.M.S is supported by an American Cancer Society Postdoctoral Fellowship (134255-PF-20-041-01-DMC). M.D. is supported in part by an NSF GRFP fellowship. We thank the Whitehead Quantitative Proteomics Core for mass spectrometry, the Whitehead Flow Cytometry Core for cell sorting, the Massachusetts General Hospital CCIB DNA core for sequencing, Océane Marescal for technical assistance, and members of the Cheeseman and Bartel labs for helpful discussions. We also thank Dave Bartel, Nicky Whiffin, and Garabet Yeretssian for helpful discussions.

## Author contributions

Conceptualization: J.L. and I.M.C.

Investigation: J.L., E.M.S., Y.T., E.M.B., E.K.

Data analysis: J.L. and M.D.

Writing: J.L. and I.M.C.

Supervision: I.M.C

Funding acquisition: I.M.C, J.L., E.M.S., M.D., Y.T.

## Methods

### Cell culture

HeLa and HEK293T cells were maintained in DMEM supplemented with 10% heat-inactivated fetal bovine serum, 2 mM L-glutamine, and 100 U/mL penicillin–streptomycin at 37 °C in a humidified incubator with 5% CO₂.

### Safe harbor (AAVS1) transgene targeting and live cell imaging

All cDNA used in this study, with the exception of constructs derived from non-human genes (mouse AKR7A2 and Orangutan ALDH9A1), was amplified from HeLa cell cDNA. The mouse AKR7A2 and orangutan ALDH9A1 cDNA was synthesized by Twist Bioscience. To prepare HeLa cell cDNA, whole cell HeLa cell RNA was reverse transcribed using the Maxima First Strand cDNA Synthesis Kit (K1641) according to the manufacturer’s protocol.

All constructs were inserted downstream of a doxycycline-inducible promoter (*41*). The donor plasmid encoded a puromycin resistance cassette, the reverse doxycycline-controlled transactivator, and the transgene of interest, flanked by homology arms targeting the AAVS1 safe-harbor locus (*72*). HeLa cells were co-transfected with 500 ng of the donor plasmid and 500 ng of pX330-AAVS1 sgRNA using Lipofectamine 2000. Forty-eight hours after transfection, cells were subjected to selection with 0.45 µg/mL puromycin for a minimum of three days.

### *In vitro* transcription, capping, polyadenylation, and purification

Plasmids containing the reporter gene with a C-terminal GFP tag were amplified via PCR with an oligo with the T7 promoter (5′-TAATACGACTCACTATAGGG-3’) to generate templates for *in vitro* transcription. The resulting PCR products were digested with DpnI to remove plasmid template, purified using EconoSpin™ All-in-1 Mini Spin Columns (Epoch Life Science, 1920-250) and eluted with RNase-free water.

For in vitro transcription, the purified PCR product was transcribed using the HiScribe T7 ARCA mRNA Kit with tailing (NEB, E2060) according to the manufacturer’s instructions. RNA purification steps were performed using the Monarch Spin RNA Cleanup Kit (NEB, T2040) as recommended by the manufacturer. Purified mRNA was eluted in nuclease-free water, quantified using a Nanodrop, aliquoted, flash-frozen with liquid nitrogen, and stored at −80 °C.

### mRNA transfections

Cells were seeded in 12-well plates and grown to ∼80% confluency prior to transfection. mRNA transfections were carried out using Lipofectamine MessengerMAX (Thermo Fisher, LMRNA008) according to the manufacturer’s protocol, using 1 µg of mRNA and 2 µL of reagent per well. Eight hours after transfection, cells were gently dissociated by incubation with PBS supplemented with 5 mM EDTA for 5 min at 37 °C. Cells were then replated onto glass-bottom 12-well imaging plates at ∼60% confluency and allowed to recover overnight.

### Live cell imaging

For dox inducible transgenes, cells were induced for 16 hours with 1 µg/mL doxycycline prior to imaging. Cells were incubated with Hoechst dye (0.1 µg/mL) for 30 min prior to imaging. Imaging was performed in a temperature-controlled chamber using a DeltaVision Ultra imaging system (Cytiva) equipped with a 60×/1.42 NA objective. Z-stack images spanning 8 µm were acquired with a step size of 0.2 µm.

### Image quantification

CellProfiler (*73*) was used to quantify the nuclear-to-cytoplasmic ratio of GFP expressed from VHL 5′ UTR reporter constructs. Nuclei were first segmented based on DNA staining using a global Otsu thresholding strategy with two classes and a minimum and maximum object size of 100–300 pixels. Cell boundaries were then defined from each nucleus using the GFP signal and the propagation method, with global Otsu thresholding and three classes.

Integrated GFP intensity was measured separately for the nuclear and cytoplasmic compartments. To restrict the analysis to transfected cells expressing the reporter, a minimum integrated GFP intensity threshold of 500 was applied to both nuclear and cytoplasmic signals.

### RNA extractions

Cells were detached using PBS supplemented with 5 mM EDTA, pelleted in DMEM, and rinsed once with PBS. The resulting pellet was lysed in 400 μL TRI reagent (Invitrogen, AM9738). Chloroform (120 μL) was added, and samples were mixed vigorously followed by centrifugation at 21,000 × g for 15 min at 4°C. The upper aqueous phase was transferred to a new tube, extracted again with an equal volume of chloroform, and centrifuged at 21,000 × g for 1 min at 4°C. For RACE, the RNA was precipitated by supplementing the aqueous phase with 300 mM NaCl and 30 μg GlycoBlue (AM9516), followed by addition of an equal volume of isopropanol and incubation at −20°C overnight. For CAGE, the RNA was precipitated with LiCl (2.5 M final) at −20°C overnight. RNA was pelleted by centrifugation at 21,000 × g for 30 min at 4°C, washed once with 70% ethanol, and resuspended in RNase-free water. RNA concentration was determined using a Nanodrop.

### 5′ Rapid Amplification of cDNA End (RACE) and sequencing

5′ RACE (*74*) was carried out using the Template Switching RT Enzyme Mix (NEB, M0466) following the supplier’s recommendations. For each reaction, 1 µg of total RNA was used as input for first-strand cDNA synthesis. Gene-specific reverse-transcription primers were first annealed to RNA by heating for 5 min at 70 °C. Reverse transcription was then performed in the presence of a template-switching oligonucleotide (5′-GCTAATCATTGCAAGCAGTGGTATCAACGCAGAGTACATrGrGrG-3′). Reactions were incubated at 42 °C for 90 min and terminated by heating to 85 °C for 5 min. Resulting cDNA was amplified by PCR using Q5 Hot Start High-Fidelity 2× Master Mix (NEB, M0494), with 5% of the reverse-transcription product used per reaction. To enhance PCR specificity, nested gene-specific primers located internal to the reverse-transcription primers were used. PCR was performed using a touchdown protocol incorporating progressively lower annealing temperatures, with initial cycles at elevated temperatures (72 °C and 70 °C) to favor target-specific products. Amplified products were resolved on 1.8% agarose gels for visualization or purified using the Zymo DNA Clean and Concentrator kit (D4004) prior to sequencing. Amplicons were sequenced using the MGH CCIB DNA Core complete amplicon sequencing service. Sequencing reads were assembled de novo, and unique 5′ end positions were quantified and displayed in 5′ RACE plots.

### 5′ UTR and transcript annotations

For the analysis of 5′ UTRs, we had to select a representative transcript per gene. We took three independent approaches:

First, we selected the most highly expressed transcript in HeLa cells based on RNA-sequencing. Given the fact that our start site profiling analysis was done in HeLa cells (*28*), the mostly highly expressed mRNA isoform can provide the strongest signal to each start codon. To do this, we reanalyzed RNA-sequencing data from control knockout HeLa cells ((*75*), 75×75 or 150bp reads) and G2/M HeLa cells ((*76*); 100×100bp reads). For Kallisto (*77*), we indexed gencode.v47.transcripts.fa (*78*) and ran Kallisto quant with default settings. We took the average of each transcript abundance, selected the most highly expressed transcript isoform per gene as the representative transcript, an approach we previously used to identify cell type specific alternative splicing events (*79*). If the transcript was not expressed based on the HeLa cell RNA-seq, we selected the longest transcript isoform. The following samples were reanalyzed: GSM9168195, GSM9168199, GSM9380031, GSM9380032, GSM9380039, GSM9380040.

Second, we used MANE select (*80*) as a community standard transcript for clinical reporting. MANE Select is a project that provides a single representative transcript per protein-coding gene that is highly reliable and consistent between the NCBI RefSeq (*81*) and Ensembl/GENCODE (*46, 78*) annotations.

Third, we selected the mRNA isoform that had the longest annotated protein coding sequence from each gene in the Gencode v25 annotations. This transcript choice is frequently used when calculating the translational efficiency of the gene and is an approach that we took in our prior studies (*28, 82*).

The numbers reported in the main text represents analysis using most highly expressed transcripts from HeLa cells (*75*).

For the assignment of mitochondrial genes, we used MitoCarta3.0 (*83*).

The sequences of AKR7A2 and ALDH9A1 was retrieved from the Ensembl database for the conservation analysis.

### Gene enrichment analysis

For quantitative measurements, such as quantitative IP-MS, we performed Gene Set Enrichment Analysis (GSEA; (*84*)) to identify pathways and processes significantly enriched in our IPs. Pre-ranked gene lists were generated based on fold-change, and enrichment scores were calculated using default parameters using the fgsea package (1.24.0) in R.

For binned or categorical analyses, such as short vs long 5′ UTR genes, we used Gorilla (*85*) with default settings to identify enriched Gene Ontology (GO) terms. REVIGO (*86*) was used to reduce redundant GO terms.

### Subcellular localization predictions

We used DeepLoc2.1 (*87*), and SignalP (*88*), webserver with default settings to predict subcellular localization.

### Cap Analysis of Gene Expression (CAGE)

For genome-wide mapping of transcription start sites, total RNA (10 µg per sample) was submitted to an external service provider for Cap Analysis of Gene Expression quality control, library preparation, and sequencing. The raw CAGE-seq reads were mapped to the human genome (GRCh38 Genome Reference Consortium Human Build 38, Dec 2013) using STAR with the default settings except with --sjdbScore 2. The resulting bam file was indexed using samtools index. Aligned reads were processed using bedtools genomecov (v2.30.0) (*89*) with the parameters −5 -bg -ibam to compute coverage restricted to the 5′ end of each aligned read, generating bedGraph files representing 5′ read end density at single-nucleotide resolution. The resulting bedGraph files were visualized in Integrative Genomics Viewer (IGV, (*90*)) to examine the distribution of read 5′ ends across genomic loci. Aligned RNA-seq reads were converted to genome-wide coverage tracks using bamCoverage (deepTools v3.5.0, (*91*)) with the parameters --binSize 1 --outFileFormat bigwig --normalizeUsing RPKM to generate normalized single-base resolution bigWig files. Enrichment around transcription start sites (based on the Gencode v25 annotations) was quantified using computeMatrix with the parameters -a 1000 -b 1000. Heatmaps were generated from the resulting matrices using plotHeatmap to visualize TSS enrichment.

To call transcription initiation sites for CAGE-Seq data, aligned reads in BAM format were processed to identify transcription start sites (TSSs) based on the 5′ ends of mapped reads, which represent capped RNA molecules. Individual CAGE transcription start sites (CTSS) were extracted from aligned reads by identifying the 5′ position of each read. CTSS positions and their associated read counts were compiled across the genome, generating a map of transcription initiation sites in HeLa cells. To normalize for library size and enable comparison across samples, CTSS expression levels were converted to tags per million (TPM) by dividing raw read counts by the total number of mapped reads and multiplying by 10^6^. Quality filtering was applied to retain only high-confidence TSSs, requiring each CTSS to meet two criteria: (1) TPM ≥ 1, and (2) minimum read count ≥ 5. Filtered CTSS positions were annotated with their nearest gene using custom start codon coordinates derived from the most highly expressed transcript based on RNA seq data in HeLa cells (cite). For each CTSS, we calculated both signed distance to the annotated start codon. Negative lengths indicate upstream of the start codon whereas positive lengths indicate downstream of the start codon. All CAGE-seq data processing was performed using custom scripts in R with the GenomicRanges (*92*) package from Bioconductor.

### GFP immunoprecipitation

To enrich GFP-tagged annotated or truncated CFD, 5 15-cm plates of cells were pelleted, washed once with cold PBS, once with 50 mM HEPES (pH 7.4), 1 mM EGTA, 1 mM MgCl₂, 300 mM KCl, and 10% glycerol, then the pellet was resuspended at a 1:1 ratio in buffer containing 50 mM HEPES (pH 7.4), 1 mM EGTA, 1 mM MgCl₂, 300 mM KCl, and 10% glycerol and snap frozen with liquid nitrogen. Cell pellets were thawed following the addition of an equal volume of lysis buffer (75 mM HEPES pH 7.4, 1.5 mM EGTA, 1.5 mM MgCl₂, 450 mM KCl, 15% glycerol, 0.075% NP-40) supplemented with EDTA-free protease inhibitors (Roche) and 1 mM PMSF. Cells were lysed by sonication, and insoluble material was removed by centrifugation at 21,000 × g for 30 min at 4°C.

Clarified lysates were incubated with 100 µL Protein A beads conjugated to rabbit anti-GFP antibodies (*93*) and rotated end-over-end for 2 hr at 4°C. Beads were washed 5 times in wash buffer (50 mM HEPES pH 7.4, 1 mM EGTA, 1 mM MgCl₂, 300 mM KCl, 10% glycerol, 0.05% NP-40, 1 mM DTT, and protease inhibitors), with each wash performed for 5 min at 4°C with rotation. Bound proteins were eluted using 3 sequential glycine elutions with 100 µL 100 mM glycine pH 2.6 per elution, and precipitated by addition of 1/5 volume trichloroacetic acid (TCA) at 4°C overnight. Precipitates were washed three times with ice-cold acetone and dried by vacuum centrifugation.

### Mass spectrometry

Protein digestion was performed using a modified S-trap workflow (Protifi). Dried TCA pellets were resuspended in S-trap lysis buffer containing 5% SDS and 50 mM TEAB (pH 8.5) and heated to 95°C for 10 min in the presence of 20 mM DTT. Cysteines were alkylated by treatment with 40 mM iodoacetamide for 30 min at room temperature, followed by acidification to a final concentration of 2.5% phosphoric acid. Six volumes of S-trap binding buffer were added, and samples were loaded onto S-trap mini columns by centrifugation at 4,000 × g. Columns were washed four times with S-trap binding buffer. On-column digestion was carried out overnight at 37°C using 1 µg trypsin in 50 mM TEAB (pH 8.5). Peptides were sequentially eluted with 20 µL 50mM TEAB at pH 8.5, then 0.2% formic acid, followed by 50% acetonitrile. The elutions were pooled and quantified using the Pierce Quantitative Fluorescent Peptide Assay (23290) then lyophilized.

Approximately 1.5 µg of digested peptides were resuspended in 50 mM TEAB (pH 8.5) and labeled using TMT10plex reagents at a 20:1 reagent-to-peptide ratio. Labeling reactions were incubated for 1 h at room temperature and quenched by addition of 0.2% hydroxylamine for 15 min. Labeled samples were pooled on ice, flash frozen, and lyophilized. Labelled peptides were fractionated using a high-pH reversed-phase peptide fractionation kit (Thermo Fisher Scientific), following the manufacturer’s recommendations for TMT-labeled samples. Fractions were pooled (1+2, 3+4, 5+6, and 7+8), flash-frozen, and dried.

Dried peptide fractions were resuspended in 0.2% formic acid to a final concentration of 250 ng/µL and analyzed on an Orbitrap Exploris 480 mass spectrometer equipped with a FAIMS Pro interface and coupled to an EASY-nLC system. Peptides were separated on a 25 cm C18 analytical column at a flow rate of 300 nL/min using a multi-step gradient of solvent B. The instrument was operated in positive ion mode with a spray voltage of 1.8 kV and an ion transfer tube temperature of 270°C. FAIMS analyses were performed using standard resolution settings with alternating compensation voltages across two injections. Full MS scans were acquired in profile mode at 120,000 resolution over an m/z range of 350–1200. The instrument used an automatic determination of maximum fill time, standard automatic gain control (AGC) target, an intensity threshold of 5×10³, with precursor selection restricted to charge states +2 to +5 and a dynamic exclusion window of 30 seconds.

Raw data files were processed using Proteome Discoverer version 2.4 (Thermo Fisher Scientific) to identify proteins and peptides. The Sequest HT (*94*) search engine was employed, using the Homo sapiens protein database (UP000005640) supplemented with EGFP sequences. The search parameters allowed for a maximum of two missed trypsin cleavage sites. Mass tolerances were set to 10 ppm for precursor ions and 0.02 Da for fragment ions. The analysis considered several post-translational modifications: dynamic phosphorylation (+79.966 Da on serine, threonine, or tyrosine), dynamic oxidation (+15.995 Da on methionine), dynamic acetylation (+42.011 Da at the N-terminus), dynamic loss of methionine (−131.04 Da at the N-terminal methionine), dynamic loss of methionine with acetylation (−89.03 Da at the N-terminal methionine), static carbamidomethylation (+57.021 Da on cysteine), static TMT6plex (+229.163 Da at any N-terminus) and TMT6plex (+229.163 Da on lysine residues). Isotope correction factors for TMT 10plex were applied according to the manufacturer’s specifications (Thermo Fisher; product number 90111, lot number VK306786). Peptide identifications were filtered using Percolator to achieve a false discovery rate (FDR) of no more than 0.01. For quantified proteins a minimum of 5 peptide spectrum matches (PSM) was applied.

### Western blotting

Cells on plates were washed once with 1x PBS. Cells were directly lysed with 1x Laemmli sample buffer (100 mM Tris pH6.8, 12.5% glycerol (v/v), 1% SDS (w/v), 0.1% bromophenol blue (w/v), 200 mM β-mercaptoethanol). For a confluent 6-well plate, cells were lysed in 200 µL, and scaled equivalently for different amounts of cells. Whole cell extracts were sonicated at 10% amplitude for 5 seconds using the Branson Digital Sonifier 450 Cell disrupter to sheer genomic DNA then boiling. Samples were separated by SDS-PAGE and transferred to PVDF or nitrocellulose. Blots were rinsed once with TBST then blocked in 5% milk at room temperature for 1 hour. Primary antibodies were diluted in 5% milk and incubated with the blot overnight at 4°C, washed for 5 minutes with TBST 4x, incubated with secondary antibody in 5% milk for 1 hour at room temperature, followed by another 4 washed with TBST, and rinsed with PBS once. Blots were imaged using an Odyssey Clx machine (LI-COR) and quantified using the Image Studio software (LI-COR). The following primary antibodies were used: anti-GFP (Roche, 11814460001), anti-alpha tubulin (AbCam, ab52866). The following secondary antibodies were used: IRDye 680RD Goat anti-Rabbit (LI-COR 92668071), IRDye 680RD Goat anti-Mouse (LI-COR 92668070), IRDye 800CW Goat anti-Rabbit (LI-COR 92632211), IRDye 800CW Goat anti-Mouse (LI-COR 92632210).

### 5′ UTR ClinVar analysis

To identify ClinVar (*51*) alleles affecting 5′ UTR length, we downloaded the ClinVar variant summary (December 26, 2025; 4,155,543 GRCh38 variants) and extracted 5′ UTR genomic intervals from GENCODE v47 MANE Select transcripts. We filtered for indels (deletions, insertions, duplications, and insertion+deletion variants) where both start and stop genomic coordinates fell within annotated 5′ UTR regions, yielding 988 variants. We then applied a minimum size threshold of ≥10 bp to focus on variants with meaningful length changes, resulting in 166 variants. Size changes were calculated using genomic coordinates for deletions and HGVS notation parsing for insertions, duplications, and complex indels. Code and analysis are available at https://github.com/cheeseman-lab/5-utr-clinvar-mutations.

---

**Table S1. 5′ UTR length and alternative N-terminal isoforms.**

**Table S2. CAGE analysis and alternative transcriptional start sites**

**Table S3. Quantitative CFD IP-MS analysis**

**Table S4. ClinVar 5′ UTR mutations.**

**Figure EV1.**
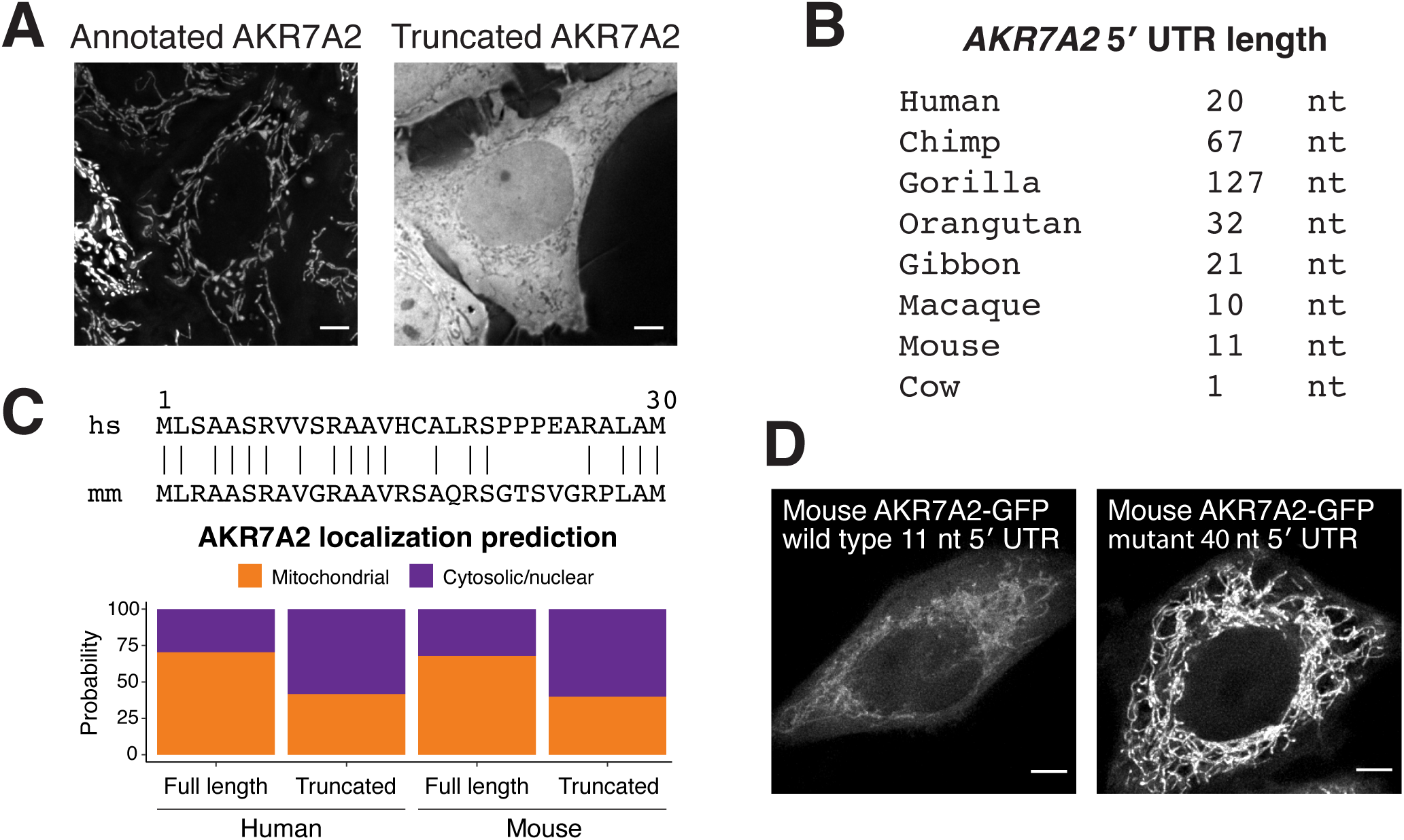
Evolutionary analysis of AKR7A2 alternative isoforms. (**A**) Live imaging of annotated human AKR7A2 and truncated AKR7A2 isoforms. (**B**) 5′ UTR of AKR7A2 across organisms. Images are not scaled equally. (**C**) Protein sequence alignment of human and mouse AKR7A2 (top). DeepLoc2.1 predictions of annotated and alternative AKR7A2 from human and mice. (**D**) Live imaging of wild type mouse AKR7A2 and a mutant AKR7A2 mRNA with a longer 5′ UTR. Scale bar indicates 5 µm.

**Figure EV2.**
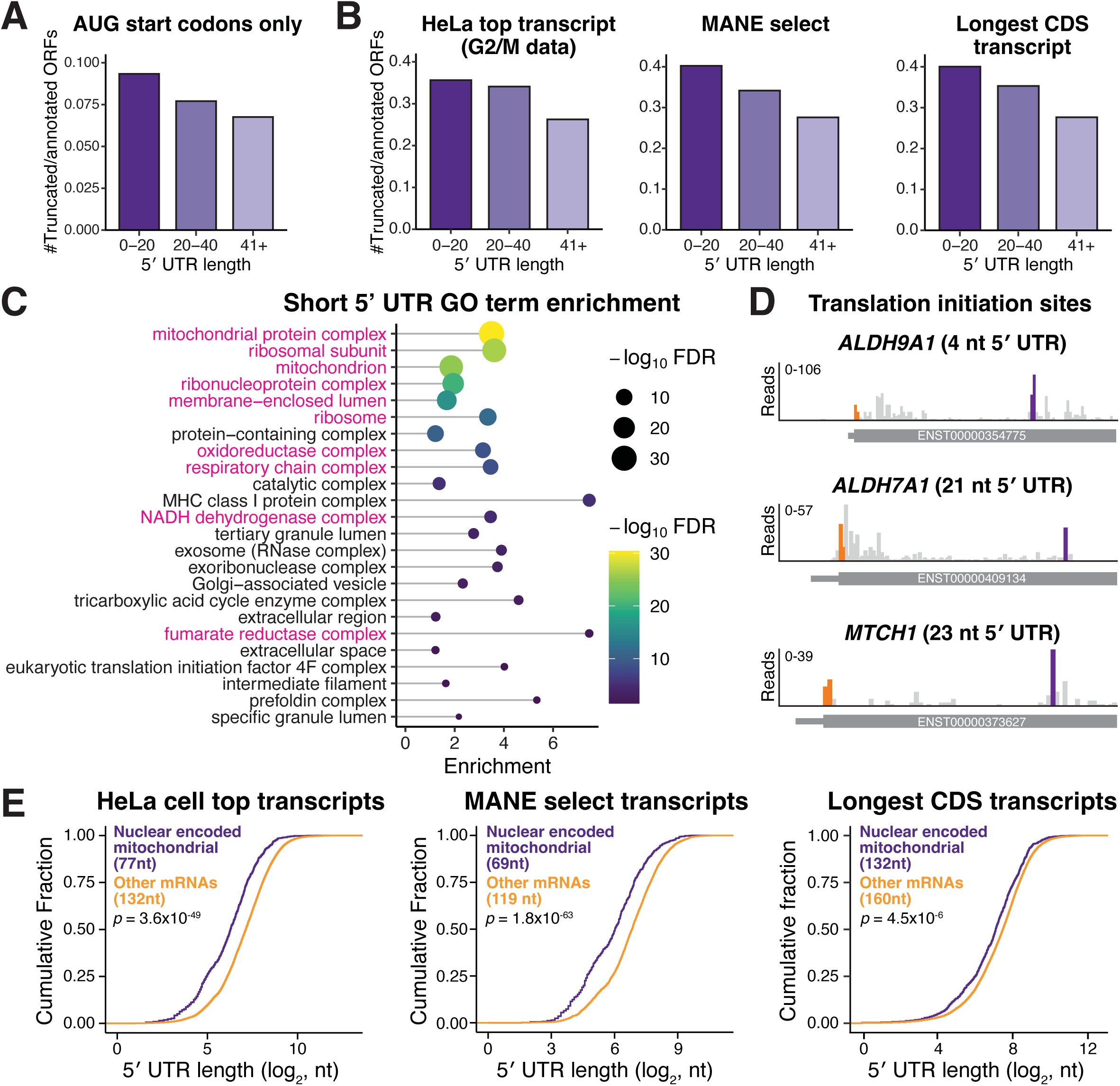
Relationship between short 5′ UTRs and N-terminally truncated protein isoforms. (**A**) Same as Fig. 2A except only AUG start codons are included in the analysis. (**B**) Same as Fig. 2A except the representative transcript isoform from which the 5′ UTR length was calculated was from different datasets (Methods). (**C**) GO term enrichment analysis of short 5′ UTR mRNAs (≤40 nt) from the MANE select transcripts (rather than highest expressed HeLa transcript in Fig. 2B) compared to transcripts with 5′ UTR lengths >40 nt. (**D**) Translation initiation site profiling traces around the start codons of indicated mRNAs. (**E**) CDF plot as described in Fig. 2C, except the representative transcript isoform for each gene was selected by different strategies (Methods). Statistics indicate Wilcoxon rank sum test.

**Figure EV3.**
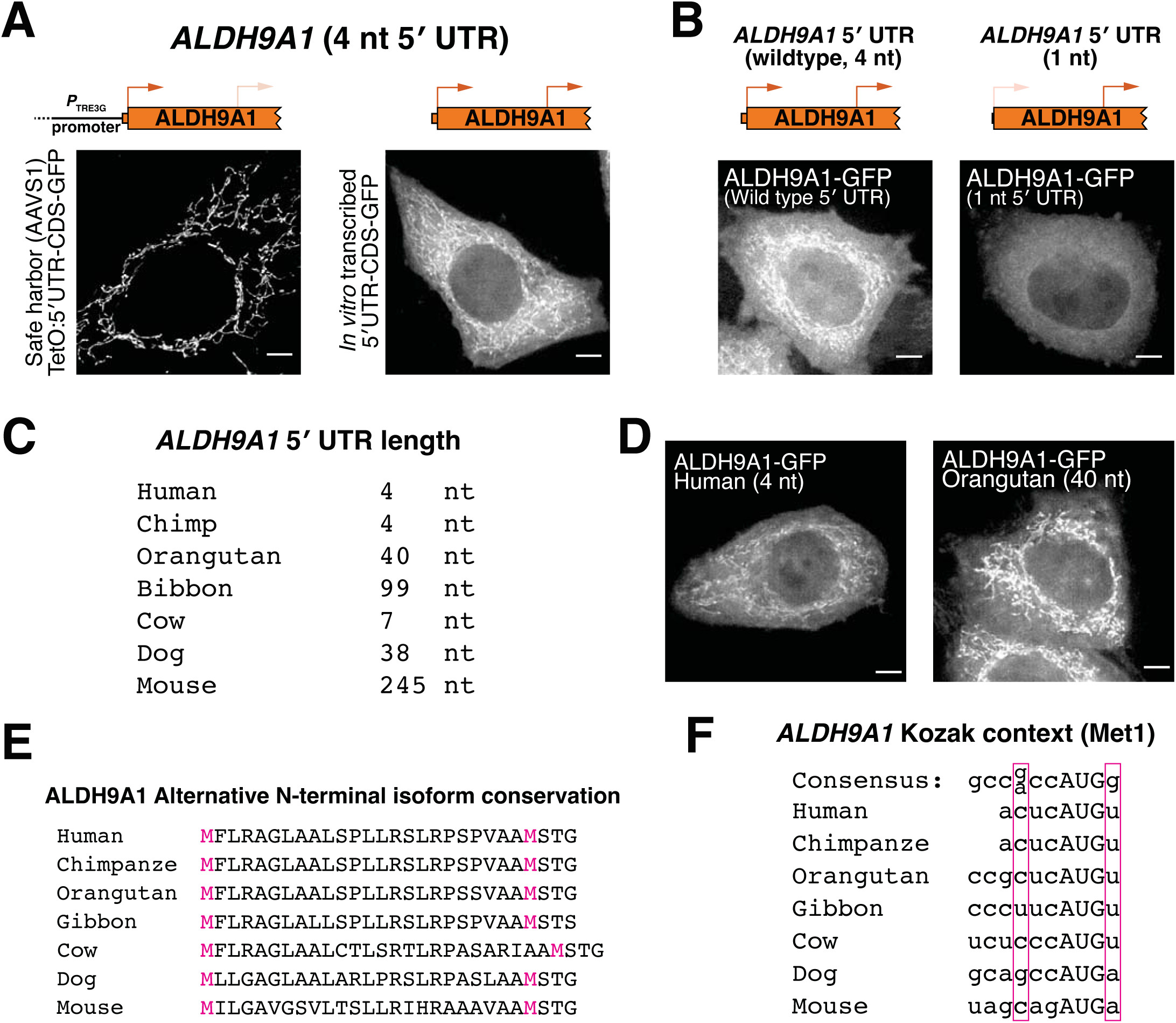
Additional analysis of ALDH9A1 start codon selection. (**A**) Live cell imaging of ALDH9A1-GFP produced from dox-inducible promoter or by transfected in vitro transcribed mRNA. (**B**) Live cell imaging of transfected in vitro transcribed ALDH9A1-GFP with the wild type (4 nt) or shorter (1 nt) 5′ UTR. (**C**) Analysis of ALDH9A1 5′ UTR lengths across organisms. (**D**) Live cell imaging of in vitro transcribed human or orangutan ALDH9A1-GFP. Scale bar indicates 5 µm. (**E**) Protein alignment for ALDH9A1 across organisms. (**F**) Conservation of weak Kozak context around the first AUG in ALDH9A1.

**Figure EV4.**
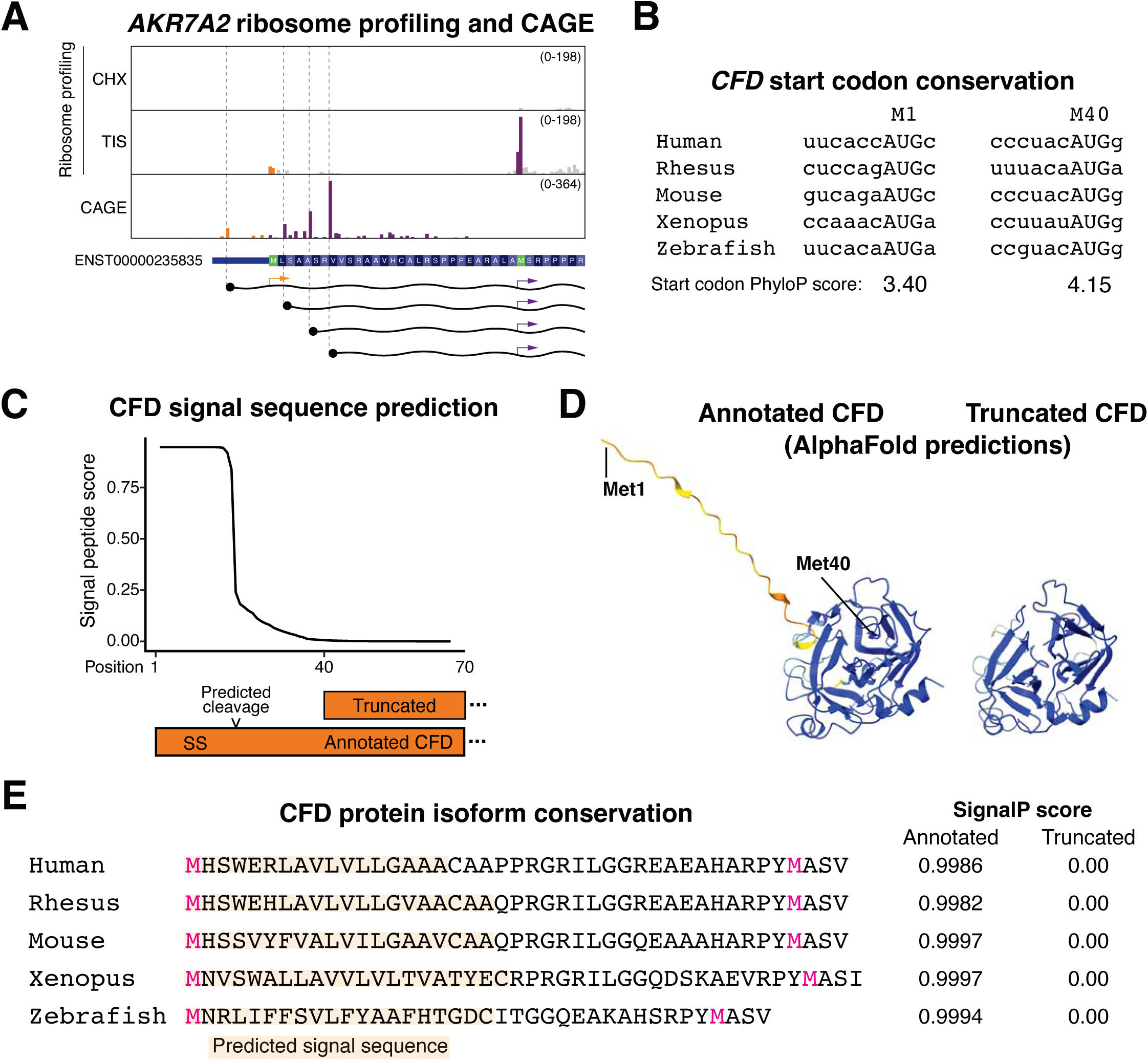
Additional analysis of alternative AKR7A2 and CFD isoform. (**A**) Read distribution traces from ribosome profiling and CAGE-seq reveals alternative promoter for AKR7A2 that removes the first AUG. (**B**) Conservation of the CFD start codon. (**C**) CFD SignalP predicts a strong N-terminal secretion signal. (**D**) AlphaFold (*95*) prediction of annotated an truncated CFD isoform. (**E**) CFD protein sequence alignment and SignalP predictions for the annotated and expected truncated CFD isoform.

**Figure EV5.**
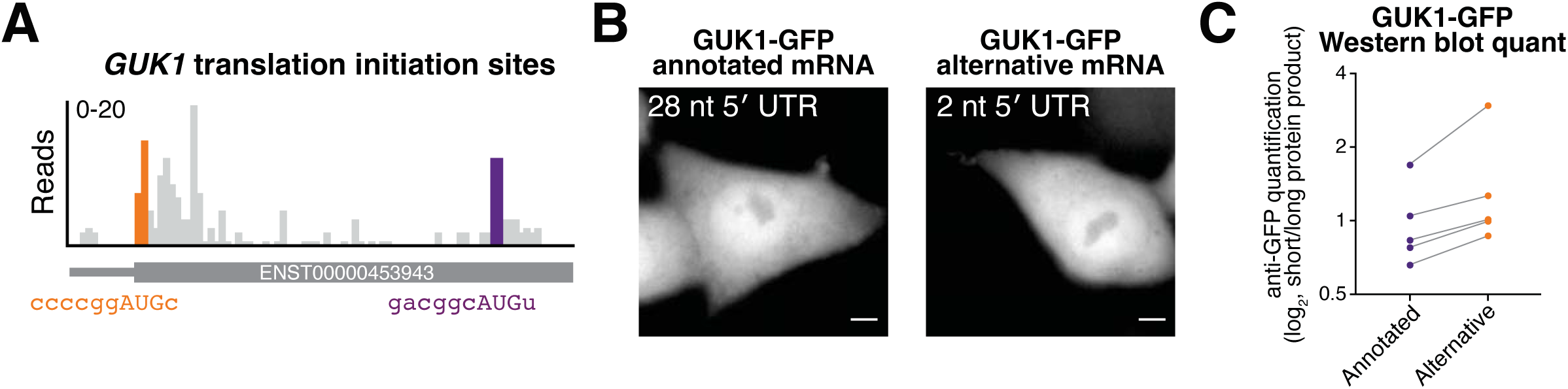
Additional analysis of GUK1 isoforms. (**A**) Translation initiation site traces around the start codon for GUK1. (**B**) Live imaging of transfected annotated or alternative GUK1 mRNA with a C-terminal GFP. Images are not scale equally. Scale bar represents 5 µm. (**C**) Quantification of Western blot from Fig. 4F. n = 5 biological replicates.

**Figure EV6.**
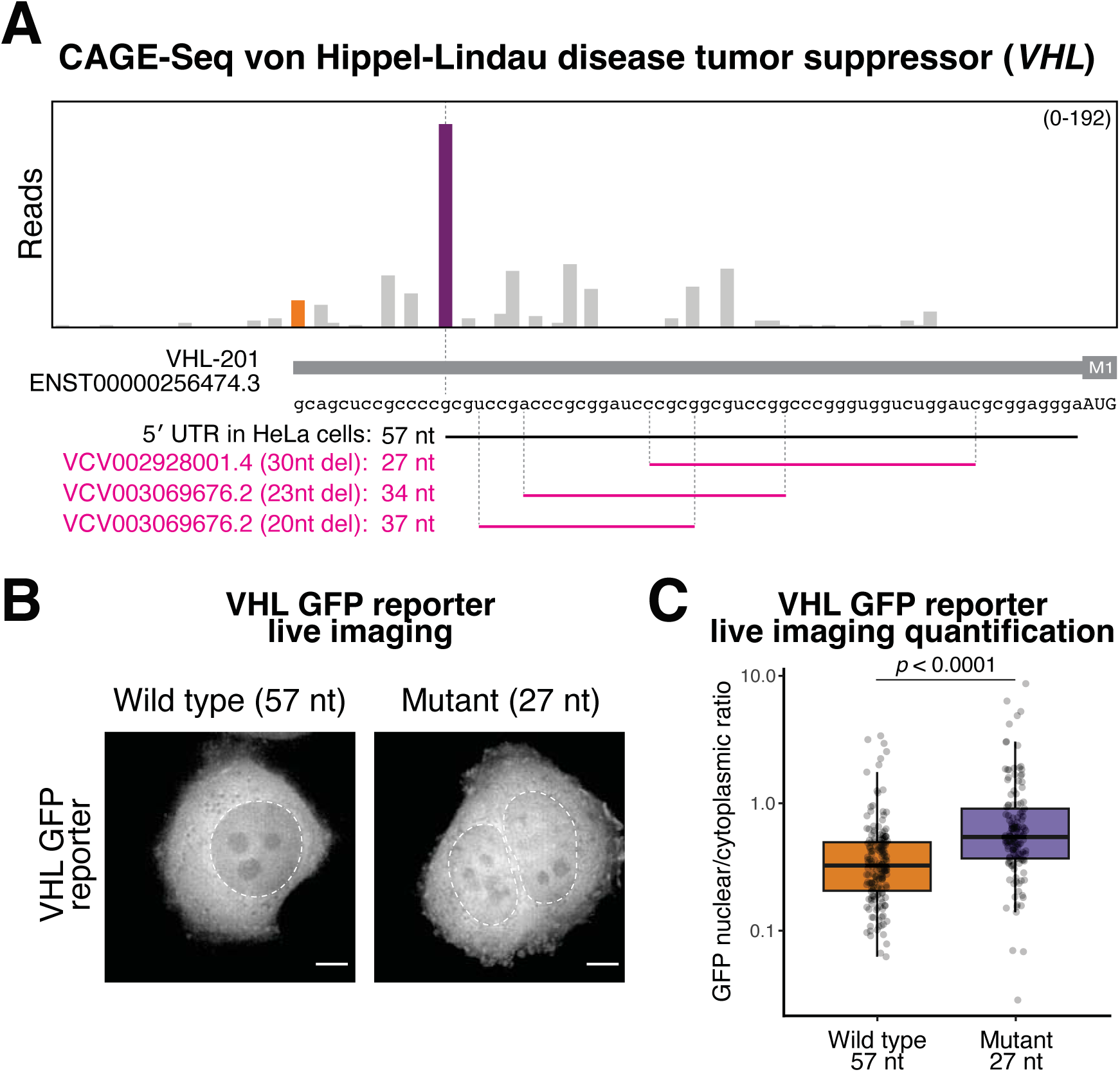
Analysis of VHL 5′ UTR pathogenic mutations. (**A**) CAGE-seq trace from HeLa cells highlighting that HeLa cells have a shorter 5′ UTR than the annotated isoform. Magenta text and lines represent ClinVar deletions. (**B**) Representative live imaging of HeLa cells transfected with the indicated in vitro transcribed reporters. Scale bar indicates 5 µm. (**C**) Quantification of nuclear vs cytoplasmic ratio of the VHL GFP reporters. Each point represents a single cell. n = 2 biological replicates.

